# Gene flow can reveal ghost lineages

**DOI:** 10.1101/2024.12.11.627931

**Authors:** Théo Tricou, Enzo Marsot, Bastien Boussau, Éric Tannier, Damien M. de Vienne

**Author notes:** Corresponding authors: Théo Tricou,; Damien de Vienne. These authors contributed equally to this work and should be considered as co-first authors (Théo Tricou, Enzo Marsot).

## Abstract

Ghost species, encompassing extinct, unknown, and unsampled taxa, vastly outnumber those typically included in phylogenetic analyses. This hidden diversity has been shown to influence the study of horizontal gene flow (e.g., introgression and horizontal gene transfer) by complicating the phylogenetic signals commonly used for their detection.

In this work, we explore the potential of horizontal gene transfer (HGT) detection methods based on phylogenies (i.e., reconciliation methods) to detect and quantify ghost diversity . We briefly outline the theoretical framework underpinning our approach. Then, using simple simulations and empirical data from Cyanobacteria, we show that HGT signals, as interpreted by reconciliation methods, can reveal the presence and phylogenetic position of ghost clades, despite the absence of genomic data for these taxa. We anticipate possible limitations and difficulties in using HGT detection to explore ghost diversity and suggest promising approaches to address or circumvent them. Altogether, this proof of concept opens new lines of research for the future: a scarce fossil record and a large proportion of unknown lineages, especially in Archaea and Bacteria, does not equate to an absence of information for evolutionary studies.

## 1 Introduction

Horizontal gene flow (HGF) refers to the transmission of genetic material across species boundaries, as opposed to the classical vertical transmission from ancestors to descendants. Processes of HGF, ranging from introgression between closely related species through hybridization to inter-domain horizontal gene transfers (HGT), play a key role in the evolution of Bacteria, Archaea, and also Eukaryotes (Thomas and Nielsen, 2005; Keeling and Palmer, 2008; Szöllősi *et al*., 2015). HGF is a major source of genetic variation, that contributed to adaptation and adaptive radiation in most plant and animal groups, including humans (reviewed in Edelman and Mallet, 2021; Reilly *et al*., 2022). It is also a primary source of innovation in most taxonomic groups, involved for instance in the spread of antibiotic resistance genes (Ochman *et al*., 2000) and in the emergence of mitochondria and chloroplasts (Andersson *et al*., 2003; Strassert *et al*., 2021).

From a phylogenetic perspective, HGF events cause discrepancies between species trees and gene trees. As such, gene flow events have long been considered a source of confusion for species tree inference because of the additional complexity they bring to phylogenetic and phylogenomic analyses (Philippe *et al*., 2017). In the last decade, however, authors have started to realize that HGF could provide valuable information for reconstructing evolutionary relationships among organisms (Abby *et al*., 2012), or help refine the dating of phylogenetic trees by determining a relative ordering of speciation events (Szöllősi *et al*., 2012; Davín *et al*., 2018; Szöllosi *et al*., 2022).

The discrepancy between gene trees and species trees can also be exploited to get insight into the evolutionary events experienced by the genes, including horizontal gene flow. Methods such as ABBA-BABA to detect introgressions (Green *et al*., 2010), and *reconciliation methods* for HGT detection (e.g., Szöllősi *et al*., 2013b, Szöllősi *et al*., 2015), all rely on the detection and quantification of these incongruences. Recently, we showed that an important aspect of gene tree-species tree discordance had been overlooked: ghost lineages, i.e. extinct, unknown, and unsampled species. These taxa, thought to represent a huge proportion of the lineages descending from the root of any phylogeny, can severely hinder the study of gene flow if they are not properly accounted for (Gogarten *et al*., 2008; Ottenburghs, 2020; Tricou *et al*., 2022a,b, 2024; Tannier *et al*., 2024).

In this work, we propose to use the signal of inferred horizontal gene transfers (HGTs) to detect and quantify ghost biodiversity along the branches of a phylogenetic tree. Convincing cases of extinct lineages or populations identified with HGF (HGT or introgression) have already been found in the past (Andam and Gogarten, 2011; Andam *et al*., 2012; Patterson *et al*., 2012; Frantz *et al*., 2015). We explore here whether the fingerprint of HGTs that are inferred within a reconstructed phylogeny of a set of sampled species – based on its reconciliation with multiple gene trees – could enable the estimation of the amount of hidden biodiversity along each branch of the species tree. To support our argument, we propose three simple *in silico* experiments and one validation on an empirical dataset that show the relationship between HGTs and ghost species. We observe that this relationship , although statistically significant, seems harder to exploit in empirical data due to complexities not included in our simulations.

We provide direction for future research to circumvent these complexities, especially using deep learning approaches. Overall, this proof of concept study opens new avenues for research on species for which no traditional fossils are (and will ever be) available.

## 2 Materials and methods

### 2.1 Rationale

Reconciliation methods, such as ALE (Szöllősi *et al*., 2015) or GeneRax (Morel *et al*., 2020), are commonly employed to detect evolutionary events like duplication, transfer, and loss (DTL). By comparing (reconciling) gene trees with a species tree, these methods can infer likely evolutionary scenarios that explain the observed phylogenetic discordance between them. An under-explored property of reconciliation approaches is that they also carry potential for exploring invisible (or ghost) diversity. We define ghost lineages in a species phylogeny as all branches and clades that descend from the root of the studied phylogeny but that have no descendants among the sampled species (like the branch termed “ghost” in Figure 1A). These ghost lineages can be either extinct or extant but unsampled. The species tree where the ghosts are absent (the species tree in Figure 1D) is called the “sampled” species tree and the lineages it contains are called “sampled” lineages. When a ghost lineage or clade transfers one or more genes to sampled lineages, peculiar incongruences between the gene trees (of the transferred genes) and species tree appear (Figure 1): the recipient species of this transfer branches in the gene tree at the original position of the ghost species (Figure 1B and C). The expected scenario inferred with a reconciliation method for such a situation is an HGT event (the **induced** transfer) originating from the branch supporting the ghost donor (the **induced** branch) and transferring to the recipient branch in the sampled tree (see Figure 1D and details in the legend ; terminology follows (Tannier *et al*., 2024)).

**Figure 1:**
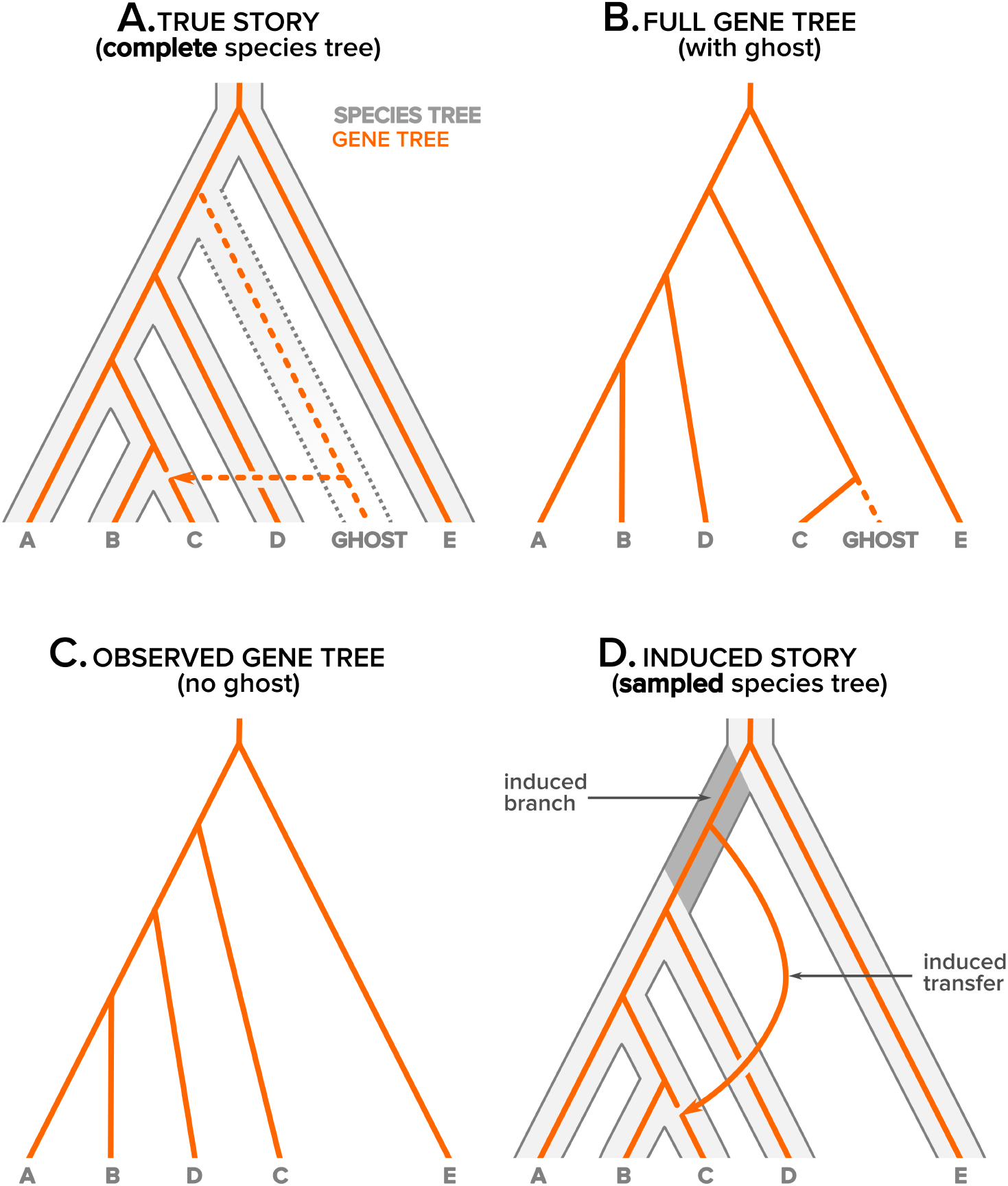
Rationale behind the use of HGT detection for getting insight into ghost species. If the true story (**A**) of the evolution of a gene (orange line) along the branches of a species tree (grey large lines) involves a transfer from a ghost branch to one of the sampled species (dashed arrow), the true gene tree (**B**) would display the recipient species (species C) as sister to the donor species (ghost). The ghost being absent from the observed gene tree (by definition), the observed gene tree will have the topology presented in (**C**). The story of the evolution of the gene in a species tree where the ghost is absent (the induced story, (**D**) should display a transfer (the induced transfer) from the branch initially carrying the ghost branch (the induced branch). Note that this induced branch corresponds to two consecutive branches of the complete species tree. If reconciliation methods correctly predict induced transfers, induced branches in the species tree should show an excess of transfers indicative of the size of the ghost clade branching on it. The **sampled** species tree in (**D**) is obtained from the **complete** species tree in (**A**) by sampling the non-ghost leaves.

We expect that if ghost lineages are abundant and transfers are frequent, many genes will follow the same transfer routes, thereby revealing the location and relative size (or transfer activity) of ghost clades. Consequently, when reconciling many gene trees with a species tree, we can expect the HGT fingerprint to show an excess of transfers originating from branches that support numerous ghost lineages (Figure 1).

In order to explore this intuition further and demonstrate the relationship between an excess of HGTs donated by a branch and the amount of ghost species it carries, we performed simulations with a simple model as well as experiments on a small biological dataset of 36 cyanobacterial genomes (retrieved from (Szöllősi *et al*., 2013a)). For the simulations, the protocol was as follows: first, we simulated species trees with ghost branches (simulated here as branches that are extant but not sampled). We then simulated genes evolving along these branches with HGT events occurring. Last, we observed the correlation between the length of a branch and the number of genes transferred from this branch, highlighting an excess of HGT events for branches bearing ghost branches. For the empirical data experiment, we performed the following test for each possible clade of the Cyanobacteria species tree. First, all species in the clade were pruned from the species tree and from all the gene trees to mimic ghost lineages. Second, all gene trees were reconciled with the species tree. Third, the number of transfers originating from each branch was recorded and compared to the number of transfers before pruning (on the full dataset). Lastly, for each branch, the difference in the number of transfers before and after pruning was computed. These differences were compared between the induced branch (initially carrying the ghost clade) and the other branches, and across the different pruning experiments.

We present below the model and the protocol used for the simulations, as well as details on the empirical data experiment.

### 2.2 General model of gene evolution under HGT

We describe the model of evolution under HGT implemented in the program Zombi (Davín *et al*., 2020), without gene duplication or loss. Let 𝒯 be a rooted, dated, binary species tree resulting from a birth-death process. This **complete** species tree (Figure 1A) contains all the species that have evolved during this process, including species that have gone extinct. These extinct species manifest as branches that stop before present time, and occur only if the death rate in the birth-death process is non-zero; otherwise the process is called a pure-birth process. After sampling a subset of extant leaves from this complete species tree, we obtain a new, smaller tree that we call the **sampled** species tree (Figure 1D). All the branches present in the complete tree but not in the sampled tree are ghost branches.

Genes evolve along the branches of the complete species tree according to two events: they either split into descendant copies during a speciation event in the species tree, or transfer with replacement (the transferred copy replaces the original copy of the gene in the recipient genome, Choi *et al*., 2012), when there is a HGT event. The topology of the gene tree is affected differently by the two events. Upon speciation events, the gene tree topology mirrors that of the species tree. Upon transfer events, the gene tree topology differs from the species tree according to a Subtree Prune and Regraft (SPR) move, which disconnects the recipient branch from its parent and reconnects it to the donor branch (see Figure 1A and B). After a succession of HGT events with the same receiver, the receiver branch will have its last donor branch as a parent, making past transfers to this branch invisible on the gene tree topology.

HGT events are distributed along the branches of the complete species tree as follows: in each infinitesimal time interval dt, any branch of the species tree donates a gene with probability λ × dt, and does not donate any gene if there is no other branch in that time interval to receive a transfer. If a gene is donated, the receiver is chosen uniformly among the other species alive at that time. We note that under this model, and in the absence of ghost species, there is a linear relationship between the expected amount of HGTs originating from a branch of the species tree and its length. In the case where there are ghost species, a transfer originating from a branch in the species tree has some chance of being received by a ghost branch, in which case this transfer will not be seen from sampled genomes. This introduces a more complicated dependency between the number of transfers originating from a branch of the sampled species tree and its branch length, involving the proportion of ghost species at the times of the transfers.

### 2.3 Simulated data generation protocol

For simplicity, we do not simulate extinct species, *i*.*e*. all ghost species in our model are unsampled species (as in Figure 1, see a justification for this choice in the discussion). We simulate, with the software Zombi (Davín *et al*., 2020), a complete species tree 𝒞 under a simple pure-birth process (without extinction)(Yule, 1925; Kendall, 1948) . Branch lengths thus represent time in arbitrary units. We then simulate the evolution of a genome of 500 genes *G*_*i*_ along the branches of this species tree. Apart from speciation, only HGTs (with replacement) are simulated, with a genome-wide transfer parameter set to 6. This corresponds to 6 transfer events expected per genome (across all 500 genes) per unit of time. Note that the number of units of times differs between the simulations, but this does not affect the expected number of transfers detected in the sampled species trees. We then remove a subset of leaves from this species tree to generate a sampled species tree 𝒮, representing the subset of species that we know. In this context, because no extinct species have been simulated, ghost species refer only to species that have not been sampled.

We perform three different simulations. In the first simulation, a complete tree with 30 leaves and with no ghost is simulated. In the second simulation, a complete tree of 40 leaves is generated, then a clade of 10 monophyletic taxa is removed to generate a clade of ghost species (30 sampled tips out of 40). In the third simulation, a complete tree of 60 leaves is generated and half of the leaves (30 out of 60) are randomly sampled, uniformly. The species trees obtained in the three simulations are displayed at the top of Figure 2. We use the software ALE (Szöllősi *et al*., 2015) to infer transfers. For each gene tree *G*_*i*_, we perform reconciliations to its associated species tree 𝒮 using the ALEml undated version of the program, under default parameters. We then summarize this information by summing the frequency of inferred transfers originating from each branch of the sampled species tree over the individual reconciliations of the *G*_*i*_ with 𝒮.

**Figure 2:**
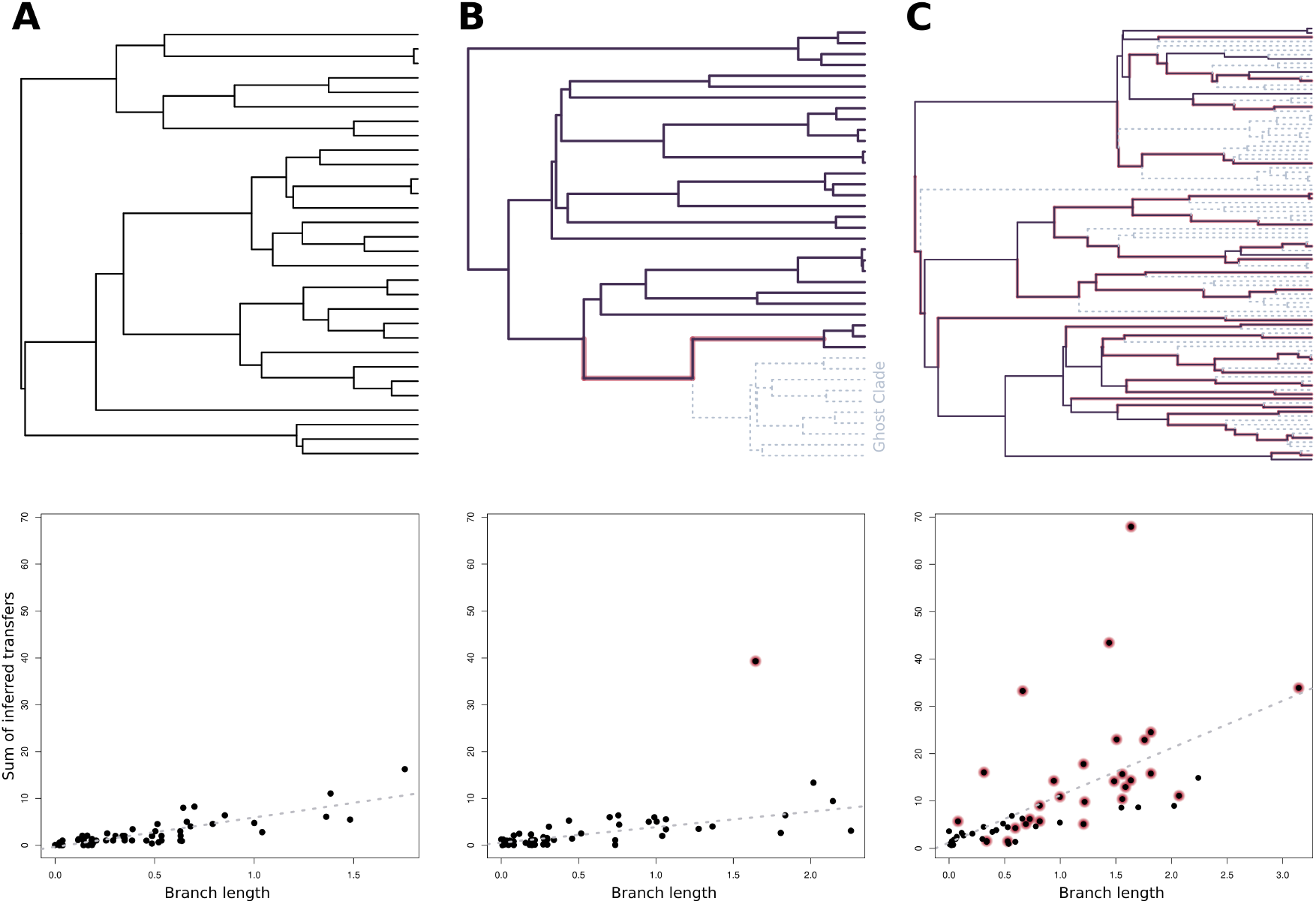
Impact of ghost lineages on the relationship between the branch lengths and the number of inferred transfers after reconciliation. (**A**). When no ghost lineages are present in the simulation, the correlation between branch lengths and inferred transfers is linear and all branches (dots) fall along the regression line. (**B**). When a single clade of 10 taxa is pruned from the species tree, the branch initially carrying the clade, the induced branch (with red background), displays an excess of inferred transfers with respect to its branch length, unlike the other branches (**C**). When 50% of the taxa (30 over 60) are randomly sampled, the linear relationship is blurred, but branches initially carrying ghost branches still display higher numbers of inferred transfers with regard to their branch lengths and this excess is proportional to the sum of ghost branch length (Figure S1). The species trees used for each simulation are displayed on top of their corresponding plot. Grey dotted branches represent the ghost lineages that were pruned from the tree prior to the reconciliations. Red haloed branches depict the branches supporting ghost lineages in the sampled species tree.

### 2.4 Test on biological data

In order to get a biological validation of the intuition presented above on the link between HGTs and ghost lineages, we set up an experiment based on a biological dataset comprising 1099 gene families from 36 cyanobacterial genomes (gene and species trees were retrieved from (Szöllősi *et al*., 2013a)). We tested whether the pruning of a clade (to mimic a ghost clade) in the tree was associated with an increase of the number of transfers originating from the corresponding induced branch, as compared to the other branches.

To do so, we started by reconciling each *full* gene tree with the *complete* species tree (nomenclature as in Figure 1) using the ALEml undated version of ALE (Szöllősi *et al*., 2015) with default parameters. All inferred events (duplications, transfers and losses) and their associated scores were recorded. We only kept - for further analysis - the genes with a moderate total sum of transfers (< 6) and low duplications (total sum of duplications < 1), to avoid working with noisy reconciliation data (a reconciliation leading to high numbers of transfers and duplications can reveal gene phylogenies that are close to random and thus carry little information). This left us with a list of 670 genes. Then, for each possible clade (including tips) of the species tree, we did the following: (i) we pruned the clade from the species tree to mimic a ghost group and produce the *sampled* species tree, (ii) we pruned all the species composing the clade from all the gene trees to produce the *observed* gene trees, and (iii) we reconciled all the *observed* gene trees with the *sampled* species tree. Note that in addition to the root for which the experiment cannot be performed, the six deepest nodes of the species tree were also excluded from the pruning experiment because transfers originating from these clades produce a signal with low specificity (i.e., many other transfers not involving the clade can produce similar unrooted gene tree topologies, thus blurring the signal, see Supplementary Material 1.2 and Figure S2). To ensure a fair comparison between the number of transfers originating from each branch before and after pruning of the ghost clade, we had to perform some actions on the list of transfers: (i) transfers with low support (ALE score < 0.5) were discarded from the analysis, (ii) transfers with the ghost clade as recipient (before pruning) were removed, as they cannot be compared with transfers after pruning ; (iii) Scores of transfers originating from the two branches that surround the ghost clade before pruning were summed as these two branches form a single one (the *induced* branch, Figure 1) after pruning.

Then, for each pruned clade, we computed the difference *d* in the number of transfers after (*tr*_*aft*_) and before (*tr*_*bef*_) the pruning, for each branch 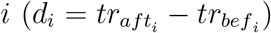. For each pruned clade, we compared *d* for the induced branch *d*_*induced*_ to *d* for the all other branches and we computed the rank of *d*_*induced*_ in the sorted list of *d* values.

## 3 Results

### 3.1 Simulations

In the first simulation (Figure 2A), where no ghosts are present, we observed a linear positive correlation (R^2^ = 0.65, p-value = 2.72e-13, dotted grey line) between the length of each branch and the summed number of transfers originating from it as inferred with the reconciliation method.

When a monophyletic group of 10 species (out of 40) was forming a ghost clade (Figure 2B), we still observed a linear positive correlation between the length of each branch and the number of transfers originating from it (R^2^ = 0.57, p-value = 1.05e-11). Short branches hardly transferred any genes and longer branches transferred more, with the regression predicting around 7 transfers for a branch of length 2.0. One branch however was an outlier with respect to this correlation with more than 40 transfers inferred when 6 were expected given its length (dot with red background). This point corresponds to the branch highlighted in red in the tree, which supported the ghost clade (left panel). This excess of transfers is due to transfers that occurred from the ghost clade to sampled lineages in the tree.

In the third simulation (Figure 2C), when a random uniform sampling of 30 tips over 60 (30 removed) was performed to create ghost lineages in the species tree, we observed a linear positive correlation between the length of a branch and the number of transfers (R^2^ = 0.37, p-value = 3.06e-07). Nodes supporting ghost lineages, highlighted in red, were scattered all across the correlation, but the excess of transfers was proportional to the sum length of all ghost branches attached to them (Figure S1).

These results give a first proof of concept that some signal extracted from the inference of HGTs could be used to estimate the amount of ghost lineages along each branch of a species tree.

### 3.2 Biological data

The pruning experiment performed on the Cyanobacteria species tree (Figure 3A) consisted in exploring the effect of removing a clade (to mimic a ghost clade as in Figure 1) on the number of transfers originating from the induced branch, *i*.*e*. the branch originally carrying the clade. All possible clades were removed and the distribution of the values of *d* for all the branches on the one hand (black boxplots in Figure 3B), and the induced branches on the other hand (red dots in Figure 3B) was computed and plotted each time.

**Figure 3:**
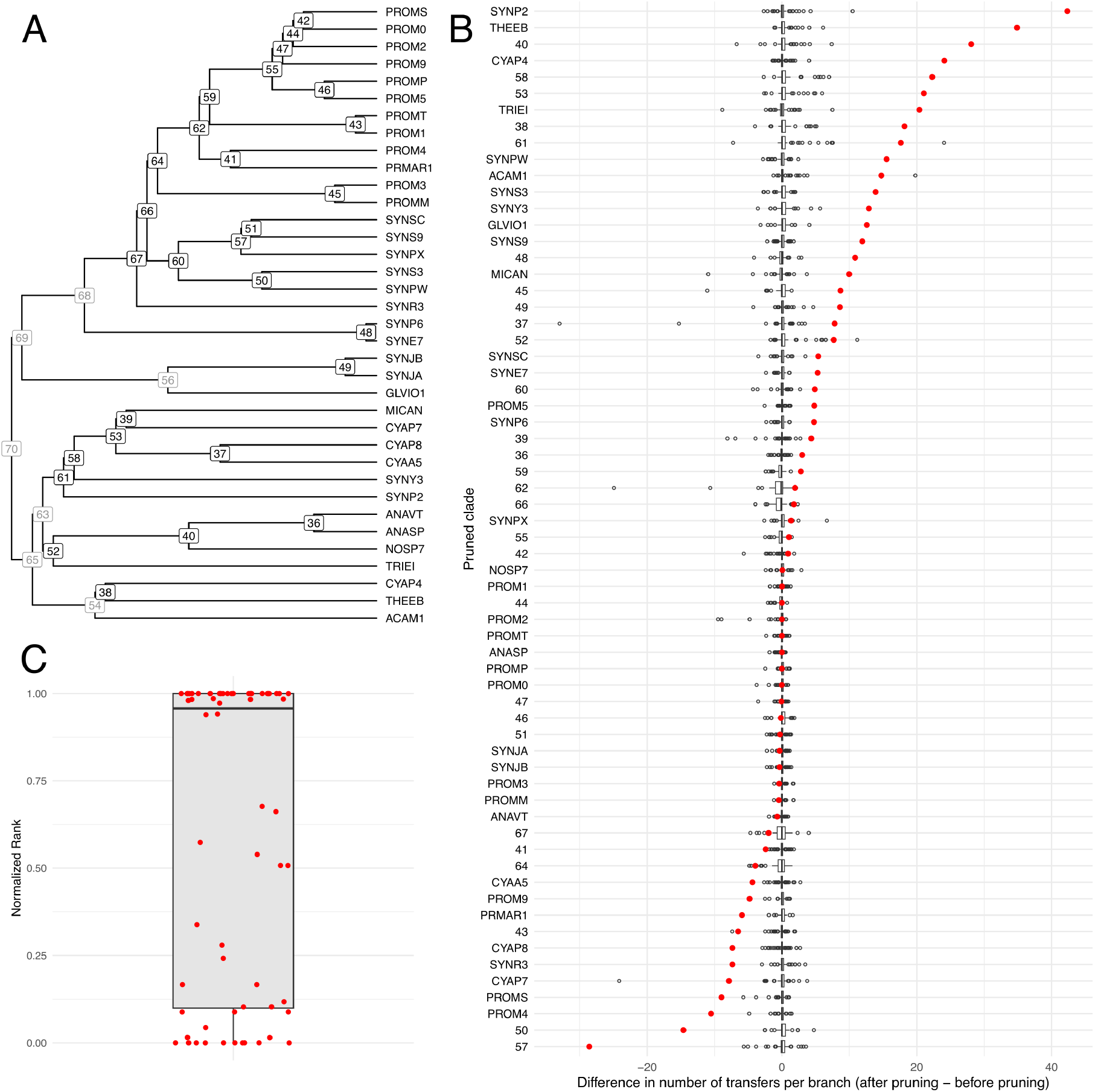
Pruning experiment on a dataset of 36 Cyanobacteria species to test the use of gene transfer detection to reveal ghost lineages. **A.** The Cyanobacteria species tree used for the experiment, retrieved from (Szöllősi *et al*., 2013a), with node and tip names as referred to in panel B. Nodes in gray were not included in the pruning experiment (see Supplementary Material 1.2 and Figure S2). **B**. Boxplots of the difference *d* (see text) in the number of transfers originating from each branch of the species tree for each pruned clade (after pruning - before pruning). Red dots represent *d*_*induced*_, the value of *d* for the induced branch. Boxplots are ordered by increasing *d*_*induced*_ values. **C**. Boxplot of the normalized ranks of the *d*_*induced*_ values in the list of *d* values. The black horizontal line represents the variance. Each dot represents an induced branch (as in panel B).

We observed that on average the difference between after pruning and before pruning was higher for the induced branches (mean = 4.5) as compared to non-induced branches (mean around 0). But we also noted that some experiments were showing an opposite trend, with a strong depletion of transfers after pruning (e.g., pruned clade 50 and 57). Nevertheless, in the sorted list of *d* values the induced branches were usually highly ranked (median of the normalized rank equal to 0.95, Figure 3C), revealing a clear tendency towards an increase of the number of transfers at the induced branch. One way to statistically test this was to compare the distribution of *d*_*induced*_ values (all the red dots of Figure 3B) to a subset of same size of *d* values for non-induced randomly sampled branches (one branch randomly sampled for each pruning experiment) using a one-sided Wilcoxon rank-sum test (Mann-Whitney U test). The sampling was repeated 1000 times, leading to 1000 p-values ranging from 0.002 and 0.07, clearly distributed below the classical 0.05 threshold for significance (Supplementary Figure S3).

Altogether, this test revealed a clear effect of pruning a clade on the variation in the number of transfers originating from the induced branch, compared to other branches, with a tendency towards increase. We also observed clade-dependent variation, including occasional decreases in transfers from the induced branch. This indicates that pruning lineages in biological datasets reshapes the gene transfer landscape predicted by reconciliation methods in a way that is, at least in some cases, hard to predict.

## 4 Discussion and conclusion

Previous studies have demonstrated that HGTs can be used to improve our understanding of the evolutionary history of sampled taxa. Here, we show on simulations and empirical data that HGTs contain information about ghost lineages, which could be used to identify extinct, unknown or unsampled lineages in species trees. This is achieved by using an existing reconciliation method, which shows that its inference of gene transfers is accurate enough to identify past donor lineages.

The toy simulations we present here illustrate how the HGT distribution in a species tree can inform on the presence, localization and size of ghost biodiversity. Many of the hypotheses underlying our reasoning may not hold true in the context of biological evolution.

First, we used a pure-birth process to generate the species tree instead of a birth-death process. While simplifying the simulations, this also eases the detection of ghost clades by increasing the strength of their signal. Indeed, unsampled ghost lineages have more time to exchange genes with sampled lineages than extinct ones do, so more transfers are susceptible to occur from the ghost clades.

Second, by assuming a constant rate of transfers, we make the assumption that the number of transfers originating from a branch is directly correlated with its length. We expect this to be violated more often than not. Third, in the simulations the recipient of a transfer is selected uniformly among the lineages alive at the time of the event. In reality, transfers are probably more likely between closely related species or species that share an ecological niche (Bolotin and Hershberg, 2017). Fourth, only simulating transfers with no duplication or loss greatly simplifies gene histories and their inference using reconciliation. Fifth, using simulation also implies that we know the true species tree topology and the true gene tree topologies. This removes the usual errors introduced by phylogenetic reconstruction done on gene sequences with limited amounts of information, which can generate spurious transfer events detection.

Our analysis of 36 Cyanobacterial genomes shows that the results obtained on our simplistic simulations hold, to a certain extent, on empirical data. Although it would remain useful to perform many additional simulations, with extinctions, with non-uniform rates of transfers, and including phylogenetic errors, these results show that transfer-based detection of ghost lineages could work in practice.

In this work, only the count of the number of transfers originating from each branch of the species tree (in simulations and in empirical data) was used to support the existence of ghost lineages. But information could come from additional types of signal, such as the identity of the recipients of the transfers. For instance, ghost lineages must be involved in cases where HGTs are inferred from a branch to one of its descendant branches. This is because the two lineages could not have been contemporaneous: a ghost lineage, branching as sister to the donor, and existing at least for some time when the recipient was alive must have received the gene from the ancestral branch, and returned it to a descendant branch. More generally, the use of branch lengths during HGT inference (neglected here, as only topologies were used for reconciliation), and the information they might convey about whether the donor and recipient branches involved in the inferred transfers were contemporaneous, could be valuable for detecting ghost lineages. New reconciliation methods that can exploit branch length information and remain efficient would be needed for this signal to be fully exploited to detect ghost lineages.

We anticipate that a way forward will be to move towards methods relying on machine learning. For instance, a neural network could be trained on simulated data to predict the amount of ghost diversity per branch. This type of approach, called simulation-based inference, has been used for phylogenetic reconstruction (Nesterenko *et al*., 2025). It could handle the complex evolutionary scenarios anticipated above, where more events of gene family evolution are simulated (duplications and losses for instance), where both species extinction and sampling are included, and where transfer rates are biased according to shared ecology or evolutionary relatedness. For such an approach to work, the species tree will need to be encoded for input in the network, for instance using Graph Neural Networks (Kipf and Welling, 2016). Similarly, the gene trees or gene alignments will also need to be encoded for input into the network. Subsequently, the network will need to interpret branch wise patterns across branches, and not a single branch at a time, to detect patterns indicative of ghost lineages. This approach is anticipated to be more effective in dealing with intricate evolutionary models compared to the widely utilized statistical frameworks of Maximum Likelihood and Bayesian inference. Although we are at the beginning of this research path and all difficulties have not been discovered, preliminary results are encouraging.

The proof of concept presented here is an important motivation for such a research program. We are confident that interesting results will come out of it. For example, large groups of species are still unknown, as shown by cultivation-independent surveys using metagenomics approaches (for instance, the CPR bacterial group, Hug *et al*., 2016). Such groups could be identified with a method relying on HGT, if transfers occurred between these ghost groups and lineages already described. The observed amount of gene sharing in Bacteria and Archaea (Beiko *et al*., 2005) is important enough that a method based on HGT could yield important discoveries.

Our work suggests that there may be ways to access extinct and unknown biodiversity in groups of species with poor fossil records and poor sampling but with a history of HGTs. More broadly, this applies to any group of species where horizontal gene flow (HGF) occurs, including eukaryotes – in which introgression is prevalent (Edelman and Mallet, 2021) – because the phylogenetic signature exploited for the detection of ghost lineages is the same regardless the type of gene flow. These new approaches could be used to learn about the evolution of biodiversity, in association with other methods that exploit the fossil record, that explore environmental metagenomic data, or that interpret the shape of reconstructed phylogenies (Morlon, 2014).

## Data Availability Statement

The data underlying this article are available in GitHub at https://github.com/SisyphusMountain/ghost_experiments.

## 1 Supplementary material

### 1.1 Relationship between ghost branch lengths and number of inferred transfers

Using the same simulation dataset as the one used for Figure 2C in the main text, we show that the excess of transfers originating from the induced branch is proportional to the size (sum of branch lengths) of the ghost clade.

**Figure S1:**
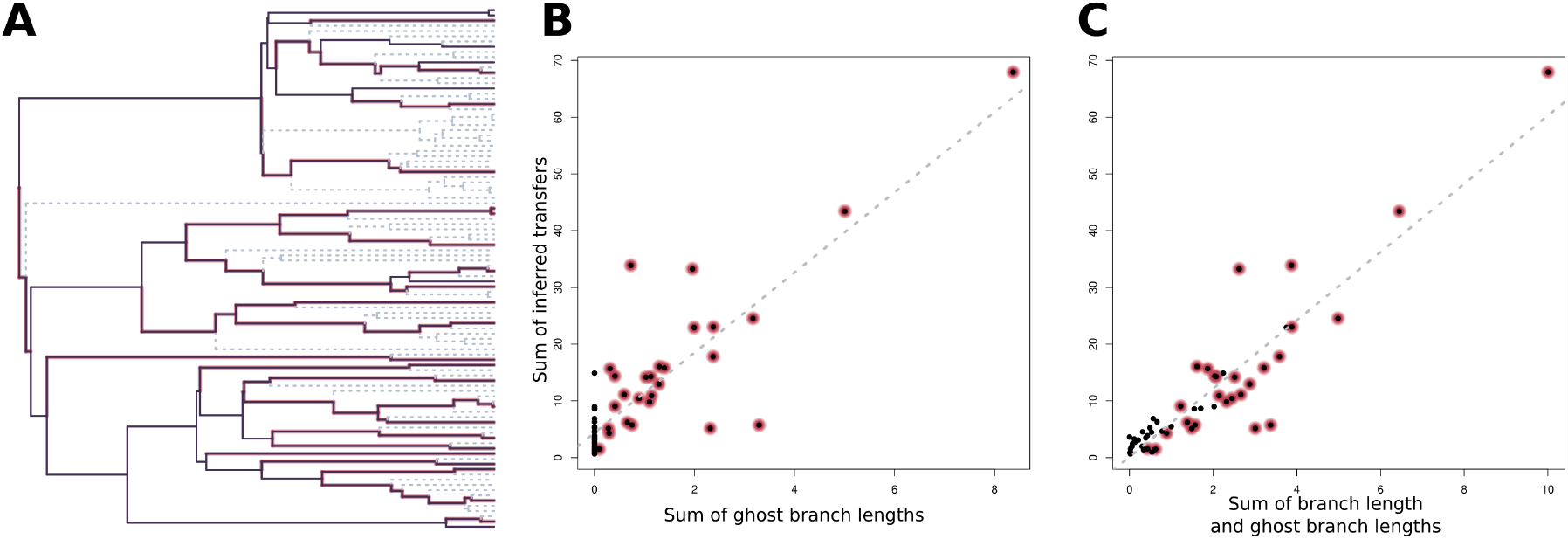
Impact of ghost lineages on the relationship between ghost branch lengths and the number of inferred transfers after reconciliation. (**A**). Species tree when 50% of the taxa (30 over 60) are randomly sampled. Grey dotted branches represent the ghost lineages that were pruned from the tree prior to the reconciliations. Red haloed branches depict the branches supporting ghost lineages in the sampled species tree. (**B**). Linear positive correlation (R^2^ = 0.7345, p-value = 2.2e-16, dotted grey line) between the length of ghost branches branching on each branch of the sampled species tree and the summed number of transfers originating from the branches in the sampled tree as inferred with the reconciliation method. (**C**). Linear positive correlation (R^2^ = 0.8492, p-value = 2.2e-16, dotted grey line) between the sum of the branch length from the sampled species tree and the length of ghost branches branching on them, and the summed number of transfers originating from the branches in the sampled tree as inferred with the reconciliation method.

### 1.2 Choice of the nodes for the pruning experiment on the biological dataset

The principle of using HGT detection for revealing ghost lineages, as depicted in Figure 1, relies on two principles. First, the expected effect of a transfer from a ghost lineage to a non-ghost lineage on the topology of the gene tree resulting from this transfer: the recipient of the transfer should branch at the position where the ghost donor was branching. Second, the expectation that the reconciliation method used for comparing the observed gene trees and the sampled species tree will indeed infer a transfer from the induced branch (see Figure 1 in main text). While the first point is obvious when we simulate (as we do here) HGTs as Subtree Pruning and Regrafting (SPR) operations, the second point is not, because transfers **not** involving the ghost clade may produce gene tree topologies similar (topologically) to those produced by the ghost transfer. In other words, some ghost clades are more likely to be detectable with transfers because the phylogenetic signature they produce is more specific than others.

Under a simple setting (*e*.*g*. a single transfer modeled as an SPR), this level of specificity can be computed for each possible ghost clade of the species tree (here the cyanobacteria species tree from (Szöllősi *et al*., 2015)) as follows:

- We simulate with SPRs all possible transfers from the ghost clade to all the other nodes (outside the ghost clade) of the tree and save the resulting unrooted tree topologies 𝒢_1_.
- We simulate with SPRs all possible transfers not involving the ghost clade and save the resulting unrooted tree topologies 𝒢_2_.
- We compute *s* the proportion of topologies in 𝒢_1_ that are found in 𝒢_2_. This represents the proportion of topologies resulting from transfers from the ghost clade that are not specific.

We computed *s* for all the possible nodes of the cyanobacteria species tree (Figure S2), except the root node (we cannot prune the full tree). We observed that the 6 deepest nodes show a value of 1, so 100% of the possible topologies resulting from transfers from these nodes (when ghosts) to other nodes of the tree can be obtained by transfers not involving the ghost clade. We removed these nodes from the pruning experiment, as they cannot be used to detect ghost clades.

**Figure S2:**
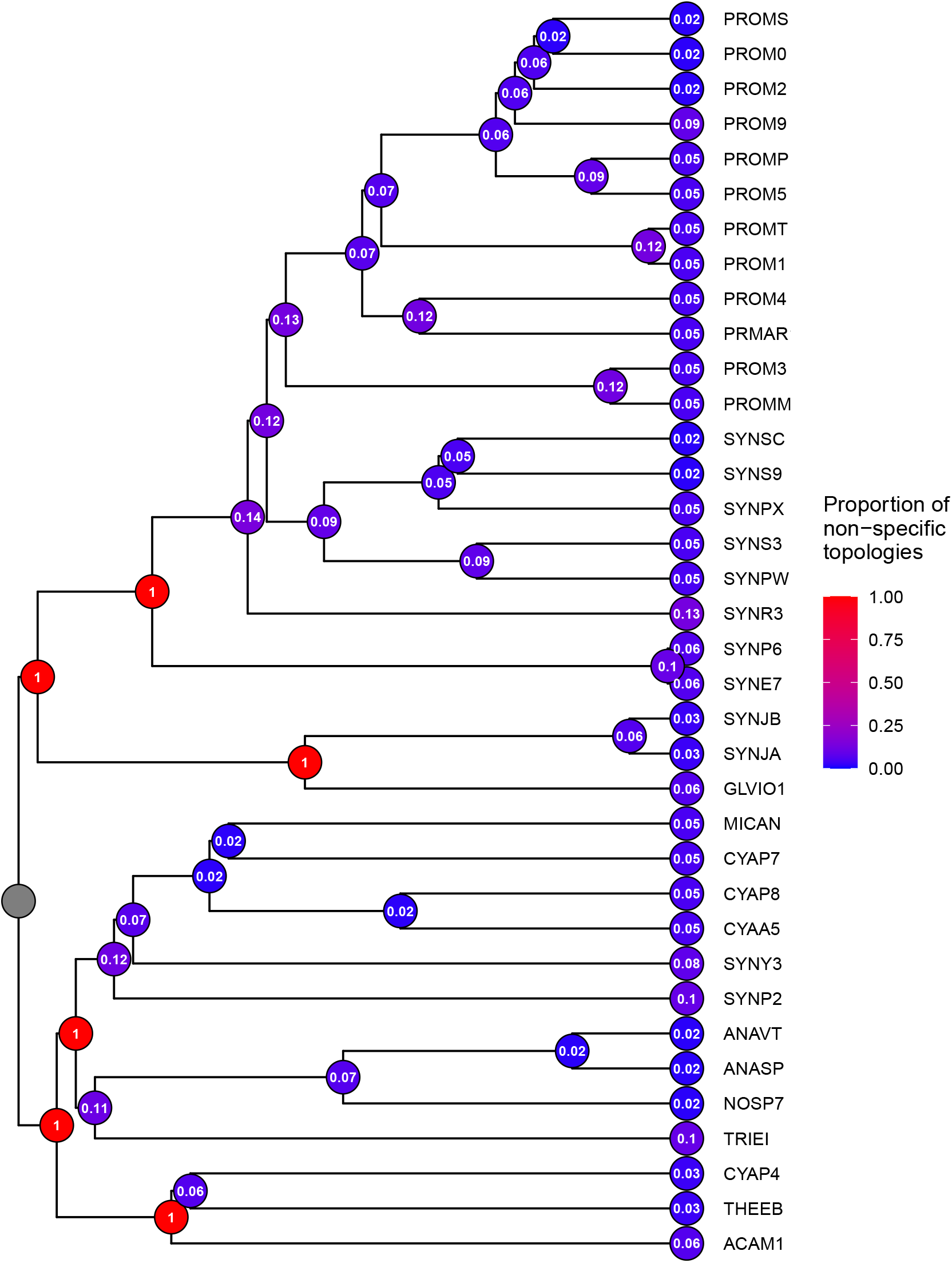
Measure of specificity of the topologies produced by SPR from each node when pruned (see text). The score represents the proportion of topologies produced by a single transfer from this node to other nodes of the tree that are not specific, *i*.*e*. that can also be obtained by transfers between other nodes of the tree. A value of 1 reveals nodes that are not suited for the pruning experiment because none of the topologies produced by transfers from them are specific. The six nodes with a value of 1 were removed from the pruning experiment.

### 1.3 Test for significance in the increase in the number of transfers originating from the induced branches

We tested with a one-sided Wilcoxon rank-sum test whether the differences in the number of transfers originating from induced branches across all pruning experiments after and before pruning was significantly higher than the distribution of differences for non-induced branches. Non-induced branches were randomly sampled (1000 times) so that the distributions had the same dimensions. One test was performed for each sampling. The distribution of the 1000 resulting p-values of the test are given in Figure S3.

**Figure S3:**
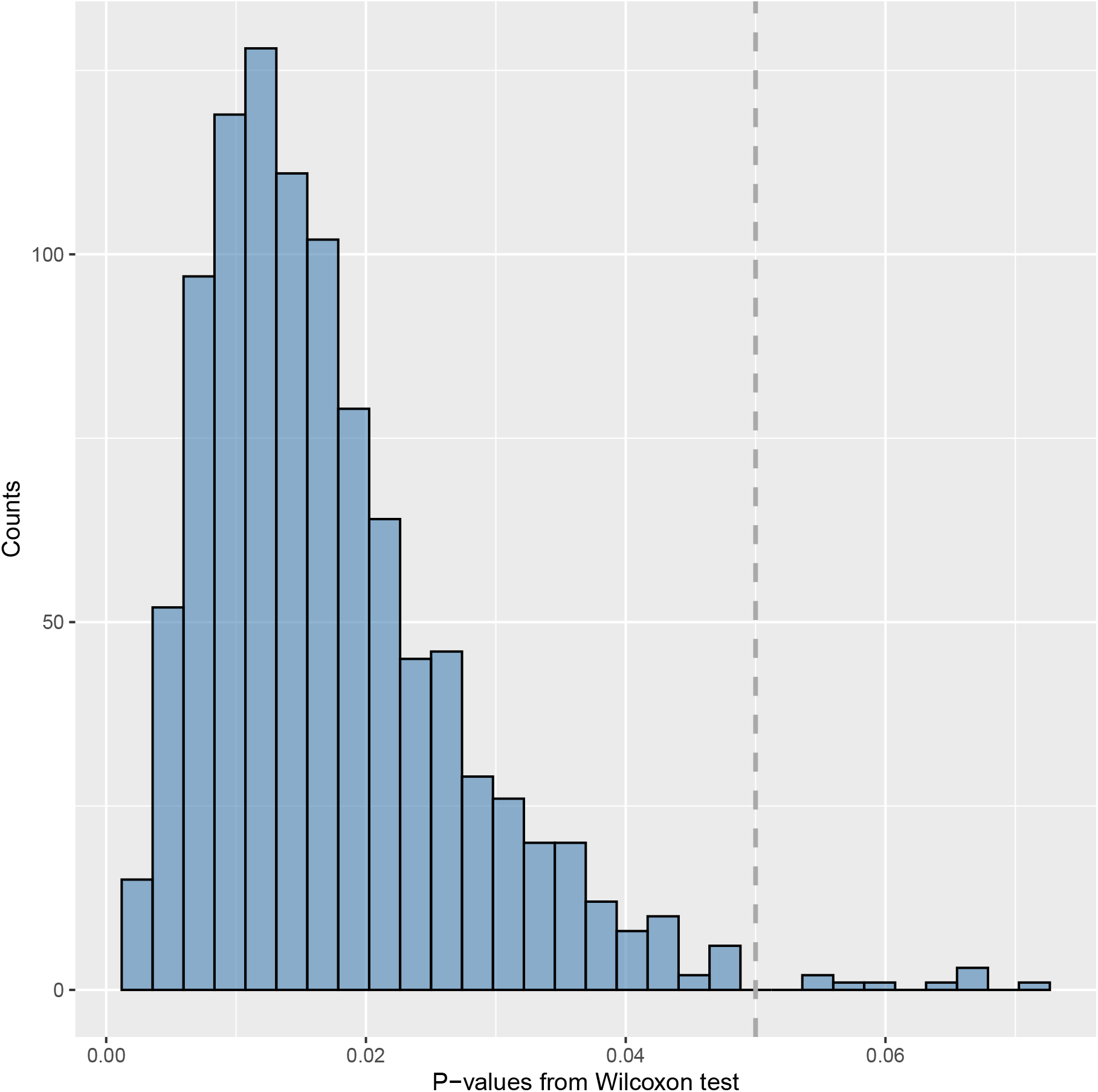
Distribution of p-values associated with the one-sided Wilcoxon rank-sum test used to explore the significance of the difference in the number of transfers between before and after pruning clades (the ghost clades), comparing the induced branches with non-induced branches. The dashed vertical gray line represents the classical significance threshold of 0.05.

